# Interpretable modelling of input-output computations in cortex

**DOI:** 10.1101/2023.08.29.555249

**Authors:** Arco Bast, Rieke Fruengel, Christiaan P.J. de Kock, Marcel Oberlaender

## Abstract

Neurons receive input from thousands of synapses, which they transform into action potentials (APs) via their complex dendrites. How the dendritic location of these inputs, their timing, strength, and presynaptic origin impact AP output remains generally unknown. Here we demonstrate how to reveal which properties of the input causally underlie AP output, and how this input-output computation is influenced by the morphology and biophysical properties of the dendrites. For this purpose, we derive analytically tractable, interpretable models of the input-output computation that layer 5 pyramidal tract neurons (L5PTs) – the major output cell type of the cerebral cortex – perform upon sensory stimulation. We find that this input-output computation is preserved across L5PTs despite morphological and biophysical diversity. We show that three features are sufficient to explain *in vivo* observed sensory responses and receptive fields of L5PTs with high accuracy: the count of active excitatory versus inhibitory synapses preceding the response, their spatial distribution on the dendrites, and the AP history. Based on this analytically tractable and interpretable description of the input-output computation, we show how to dissect the contributions of different input populations in thalamus and cortex to sensory responses of L5PTs. Thus, our approach provides a roadmap for revealing cellular input-output computations across different *in vivo* conditions.

**Author Summary:** Revealing how synaptic inputs drive action potential output is one of the major challenges in neuroscience research. An increasing number of approaches therefore seek to combine detailed measurements at synaptic, cellular and network scales into biologically realistic brain models. Indeed, such models have started to make empirically testable predictions about the inputs that underlie *in vivo* observed activity patterns. However, the enormous complexity of these models generally prevents the derivation of interpretable descriptions that explain how neurons transform synaptic input into action potential output, and how these input-output computations depend on synaptic, cellular and network properties. Here we introduce an approach to reveal input-output computations that neurons in the cerebral cortex perform upon sensory stimulation. We reduce a realistic multi-scale cortex model to the minimal description that accounts for *in vivo* observed responses. Thereby, we identify the input-output computation that these cortical neurons perform under this *in vivo* condition, and we show that this computation is preserved across neurons despite morphological and biophysical diversity. Our approach provides analytically tractable and interpretable descriptions of neuronal input-output computations during specific *in vivo* conditions.

## Introduction

Dissecting how neurons transform synaptic input into action potential (AP) output is a prerequisite for understanding the neurobiological implementation of brain functions. In the cerebral cortex, pyramidal neurons receive synaptic input from thousands of neurons along their morphologically extensive and biophysically complex dendritic trees, which they then transform into APs. When and where on the dendrite synapses are active, the ‘spatiotemporal input patterns’, is highly variable from cell to cell, and even from trial to trial within the same experimental condition [1–5]. As a result, which features of these spatiotemporal synaptic input patterns determine AP output, how this transformation depends on variability in morphological and biophysical properties, and which mathematical operation the neuron performs – i.e. what is the ‘input-output computation’ – remains unclear.

Here we derive the input-output computation that layer 5 pyramidal tract neurons (L5PTs) in the vibrissae-related part of the rat primary somatosensory cortex (vS1) – the barrel cortex [6] – perform to transform sensory-evoked synaptic input into AP output. Along their extensive and biophysically complex dendrites, these major cortical output cells receive synaptic input patterns from different thalamocortical, intracortical and top-down corticocortical pathways (reviewed in [7]). In response to sensory stimulation, L5PTs generate fast and reliable AP output with receptive fields that show large cell-to-cell variability and which are broader than those of any other cell type in the same cortical column [8].

How do these AP responses arise from the interplay of morphological, biophysical and input properties? How does the transformation from synaptic input to AP output depend on variable morphological and biophysical properties of the dendrite? Joint measurement of sensory-evoked synaptic input patterns (i.e. precise time points, dendritic locations, and origins of all active synapses), the morphology of the dendrite, its biophysical properties (which ion channels are active when and where on the dendrite) and AP output would be ideal to answer these questions, but are difficult to assess experimentally. Therefore, we have previously reported and validated a multi-scale model of the rat barrel cortex, which provides realistic estimates for the number and locations of synaptic inputs that impinge onto L5PT dendrites upon sensory stimulation, and how these dendrites transform these inputs into AP output [9–11]. Simulations of the multi-scale model captured the fast and broadly tuned sensory responses of L5PTs *in vivo*, and thereby provided concrete predictions about the underlying cellular and circuit mechanisms, which we have tested experimentally [10]. Thus, the multi-scale model sets the stage to investigate which input-output computation L5PTs perform upon sensory stimulation, and how this computation depends on the morphological and biophysical properties of the dendrites.

Here we address these questions by introducing an approach that seeks to reduce the multi-scale simulations of sensory-evoked cortical output into analytically tractable, interpretable models that capture the input-output computations during these experimental conditions, while maintaining trial-to-trial and cell-to-cell variability. The reduction revealed that the input-output computation of L5PTs can be explained by three features: the count of active synapses in a time window, their soma-distance-dependent spatial distribution on the dendrite, and the time since the previous AP of the L5PT. We find that the input-output computation is preserved across morphologically and biophysically diverse L5PTs. Based on these reduced models, we find that mostly variability in spatiotemporal input patterns and to a lesser degree in biophysical properties or morphology account for variability in AP output and receptive field shape. Furthermore, based on this input-output computation, we can dissect how thalamocortical and different intracortical input pathways contribute to sensory-evoked responses of L5PTs.

## Results

We extended our previously reported multi-scale modelling approach [10] to account for cell-to-cell and trial-to-trial variability. For this purpose, we repeated our simulations of L5PT responses to passive deflections of nine different whiskers, the somatotopically aligned / principal whisker (PW) or any of the eight surrounding whiskers (SW) (**Fig. 1A**). Whereas our previous simulations were performed on the morphology of a single *in vivo* recorded L5PT, we now performed simulations for five morphologically diverse L5PTs (**Fig. 1B**) for which we had measured highly variable receptive fields *in vivo* (**Fig. 1C**). We embedded these morphologies into the network model of the barrel cortex to provide realistic estimates for which neurons in thalamus and barrel cortex (**Fig. 1D**) could provide input to these *in vivo* recorded L5PTs, and where along their dendrites these inputs could occur (**Fig. 1E**). The network model thereby provided the spatial distributions of synaptic input patterns to L5PTs from different types of excitatory and inhibitory neurons across all layers of the barrel cortex, and from the ventral posterior medial nucleus (VPM) – the primary thalamus of the whisker system. We embedded each morphology at eighty-one locations (**Fig. S1**) in and around the C2 barrel column of the network model, and activated presynaptic neurons in the network model according to cell type- and layer-specific experimental recording data from passive single whisker deflections (**Fig. S2**). Thereby, for each morphology, each location of the network embedding, and each configuration of active neurons in the network, we generate a unique but empirically well-constrained spatiotemporal synaptic input pattern to L5PT dendrites (**Fig. 1F**). Variations in dendrite morphology and network embedding hence account for cell-to-cell variability in input in the model, and variations in network activity account for trial-to-trial variability. The hence predicted spatiotemporal synaptic input patterns (**Fig. 1G**) are remarkably consistent with those observed empirically via dendritic spine imaging [5].

**Fig 1:**
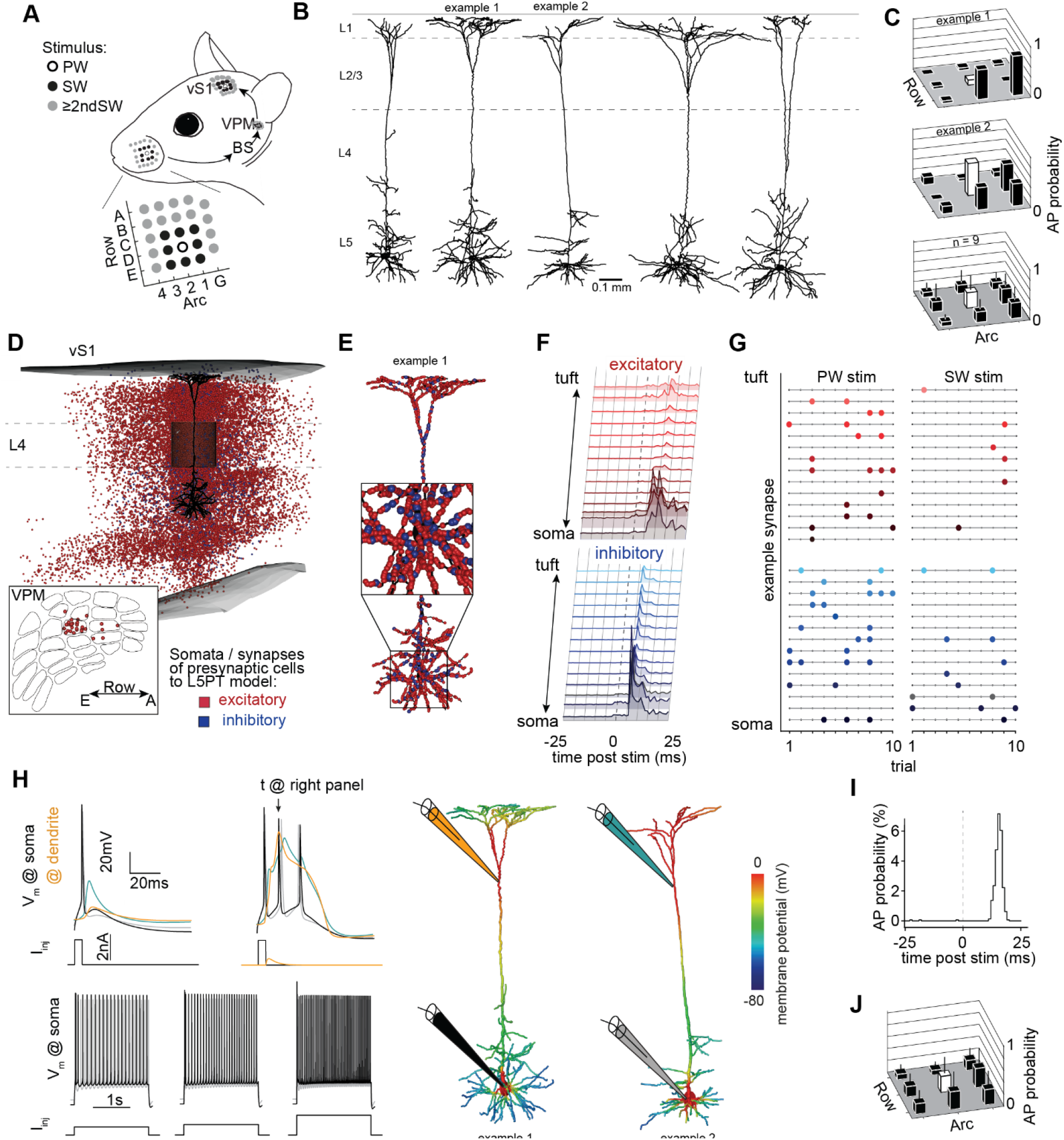
Biophysically detailed multi-scale model of whisker touch evoked responses in cortical pyramidal tract neurons. **A:** Sensory-evoked signal flow: stimuli of single whiskers (which are arranged in ‘arcs’ and ‘rows’ on the animal’s snout) are transmitted to the brainstem (BS), from there to the VPM thalamus, and from there to the primary sensory cortex of the vibrissal system (vS1). This pathway is somatotopically organized, with barreloids in VPM and barrels in vS1 corresponding to the respective whiskers. **B:** *In vivo* labeled L5PT dendrite morphologies used in this study. **C:** Corresponding receptive fields to passive single whisker touch, measured *in vivo* (upper panels). Average receptive field across 9 *in vivo* recorded L5PTs (lower panel). Error bars are std. **D:** Network model of rat vS1 and VPM provides anatomically realistic estimates of which neurons are connected to a L5PT embedded into the network. In this study, the simulated neurons are located in the C2 column of vS1, thus we refer to the somatotopically aligned C2 whisker as the ‘principal’ whisker, and the adjacent whiskers as surround whiskers. Red and blue markers denote soma locations of presynaptic excitatory and inhibitory somata, respectively. **E:** Synapse distribution originating from the neurons shown in Panel B. **F:** Spatiotemporal input pattern to L5PT: combining the anatomical constraints with empirical measurements of the activity of different presynaptic populations (Fig. S1) provides spatiotemporal input patterns that the L5PT can receives after sensory stimulation. **G:** Trial-to-trial activity of example synapses matching the soma distance from panel F for a principal whisker (C2) and surround whisker (D2) stimulus. **H:** Biophysically detailed multi-compartmental L5PT models reproduce the cell type’s characteristic electrophysiology (left panel), i.e. back propagation of APs (upper left), dendritic Ca-APs and somatic burst firing (upper right), as well as regular firing properties (lower row). Right panel: biophysically detailed neuron morphologies at the moment of a dendritic Ca AP. **I:** Simulated response to principal whisker touch. **J:** Simulated receptive fields across morphologically and biophysically diverse L5PT multi-compartmental models across 81 network embeddings capture broad and heterogeneous receptive fields.

To simulate how such spatiotemporal input patterns are transformed into AP output, we converted the L5PTs into biophysically detailed multi-compartmental models (n=7 from 5 morphologies), which capture their characteristic electrophysiological properties [12], including backpropagating APs, dendritic calcium APs and responses to step current injections (**Fig. 1H**). Each multi-compartmental model used largely different biophysical parameters to achieve these electrophysiological properties, reflecting different densities of active conductances in different subcellular compartments (**Table S1**). Thus, the embedding of these diverse multi-compartmental models into the network model allowed us to investigate how cell-to-cell and trial-to-trial variability in synaptic input, in conjunction with cell-to-cell variability in morphological and biophysical properties of the dendrites, impact sensory responses of L5PTs. Simulations of each of these multi-scale model configurations predicted the characteristic fast responses (**Fig. 1I**) and broad receptive fields of L5PTs (**Fig. 1J**). Moreover, the variability of *in silico* responses across multi-scale model configurations matched the cell-to-cell variability observed *in vivo* across L5PTs (**Fig. 1J**). Thus, these multi-scale model configurations set the stage to investigate which features of synaptic input patterns determine AP output, and how this transformation depends on cell-to-cell variability in morphological and biophysical properties of the dendrites.

### Interpretable input-output computation of L5PTs

We developed our reduction approach by using one of the multi-compartmental models, whose morphology is shown as example 1 in Figure 1B. For each of its eighty-one network embeddings, we simulated its responses to passive whisker deflections of the PW and the eight SWs, respectively. Within this simulation data, we searched for features in the synaptic input that are most predictive for the generation of an AP at a given millisecond -the ‘prediction time point’ (**Fig. 2A**, methods). For this purpose, we grouped synapses according to their activation time points (1ms bins), by their path-length distance to the soma (50µm bins), and depending on whether they are excitatory or inhibitory. We found that a weighted count of active excitatory versus inhibitory synapses, where the contribution of each synapse is weighted depending on its soma distance and activation time point, can predict AP output. We therefore determined the spatial and temporal weights (the “spatiotemporal filter”) that maximize the AP output prediction accuracy (methods). According to these weights (**Fig. 2B**), the contribution of both excitatory and inhibitory inputs of a synapse to AP output decays gradually with soma distance, reaching approximately zero at 500µm distance. Thereby, primarily proximal (i.e., basal and apical oblique) dendrites contribute to AP output upon sensory stimulation. In the temporal dimension, the contribution of excitatory and inhibitory synapses has a time course resembling excitatory / inhibitory postsynaptic potentials (EPSPs / IPSPs). Excitatory and inhibitory synapses contribute the most to AP output if they were active around 4ms and 9ms before the prediction time point, respectively. These observations were robust, independent of the method used for estimating the weights (methods, **Fig. S3**). The probability to observe a sensory-evoked AP in the multi-scale simulations increased with this weighted count of active synapses (**Fig. 2C**), which we in the following refer to as ‘weighted net input’ (WNI). The WNI predicted AP output with high accuracy (AUROC 0.949).

**Figure 2:**
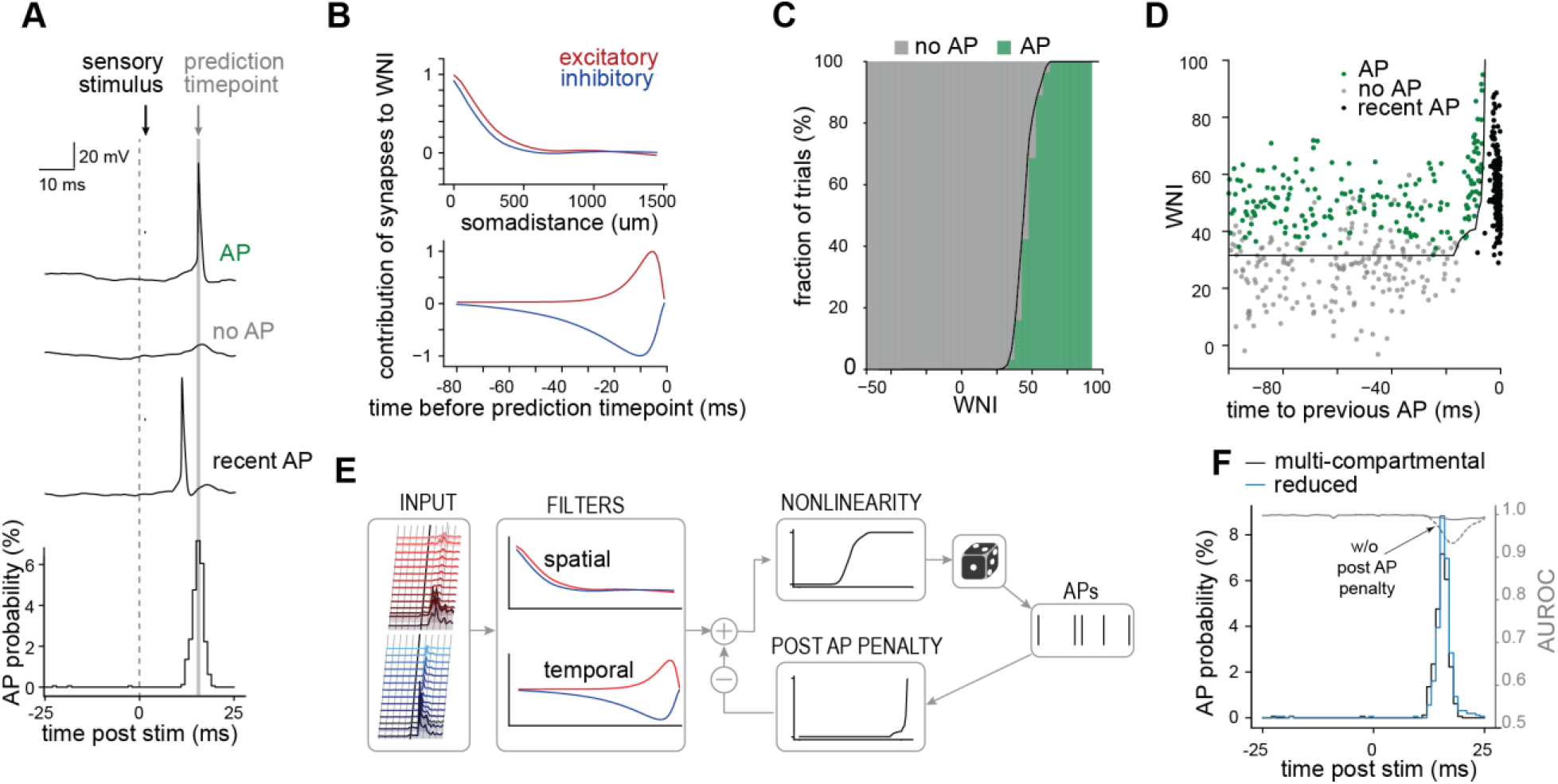
Interpretable input-output computation of L5PTs. **A:** Exemplary responses of the multi-compartmental model with respect to the prediction timepoint (the timepoint for which the occurrence of an AP is to be predicted) for the three relevant response categories ‘AP’, ‘no AP’, and ‘recent AP’ (AP was elicited shortly before the prediction timepoint). **B:** Spatiotemporal input filter that best separates AP and no AP trials assigns strong weight to proximal synapses (top) active in a short time window before the prediction timepoint (bottom). **C:** Weighted net input – the input filtered by the spatiotemporal filter – separates AP and no AP trials, but not ‘recent AP’ trials, which can be distinguished based on a second measure, ‘time to previous AP’. **D:** nonlinear relationship between WNI and AP probability. WNI represents the ‘drive’ a neuron receives; the higher the WNI the higher the probability an AP will be generated. **E:** Reduced model structure. APs are generated stochastically based on the AP probability (output of the nonlinearity). If an AP is generated, subsequent APs become less likely due to the post AP penalty, which is subtracted from the WNI. This reduced model is interpretable as it allows to directly relate AP output to synaptic input and previously generated APs. **F:** The reduced model’s responses match the biophysically detailed model across many trials (close PSTH match) and on the single trial level (high AUROC score across all timepoints). Without the post AP penalty, the AUROC score drops during the sensory-evoked response.

We revisited trials which were misclassified by the WNI and found that errors mostly occur if there was a recent AP a few milliseconds before the prediction time point. We found that the multi-compartmental model is less excitable shortly after an AP, reflecting time constants of the involved ion channels, and thus the chance of eliciting an AP is low even if the WNI is high. We therefore distinguished three categories: no AP, AP and recent AP (**Fig. 2A**). We show that these three categories can be separated based on the WNI and time to previous AP (**Fig. 2D**). To include this AP history-dependent relationship, we determined the separating line of response trials from the others (black line, methods) and incorporated it as a ‘post AP penalty’, i.e., we first computed the WNI as before and then subtracted a penalty value depending on the time since the previous AP. This penalized WNI predicted AP output with even higher accuracy (AUROC 0.990) than WNI alone. Thus, these three features, the spatial and temporal component of the WNI and the post AP penalty are sufficient to predict AP output from synaptic input.

These three features allowed us to describe the input-output computation of the biophysically detailed multi-compartmental model by an analytically tractable and interpretable model. For this purpose, we assembled post AP penalty and the nonlinear relationship between WNI and AP probability into a generalized linear model (GLM, **Fig. 2E**). We applied the GLM to predict AP output at different time points ranging from 25ms before to 25ms after sensory stimulation. Notably, even though the GLM was developed to predict the peak of AP responses, it also maintained a high AUROC score before and after this time point (**Fig. 2F**). Thus, the GLM accurately predicts APs throughout the whole time interval, and the post-stimulus time histogram (PSTH) predicted by the GLM hence matched the full PSTH simulated with the multi-compartmental model (**Fig. 2F**).

We investigated how the performance of the GLM depends on the three features it is based on. Including the AP history (i.e., post AP penalty) into the GLM is necessary to capture the input-output computation upon sensory stimulation, otherwise the AUROC score drops during the peak response (**Fig. 2F**, dashed line). Furthermore, when we simplified the GLM to neglect the spatial dimension, such that the weighted count only considers the time point of activation, but not the soma distance, the AUROC score decreased (**Fig. S3A, S3E**). It was also insufficient to incorporate the spatial dimension categorically by only distinguishing a proximal and a distal compartment (**Fig. S3B, S3E**). Similarly, when we replaced the EPSP/IPSP like shape of the temporal weights with rectangular shapes that reflect a time window with a defined start and end, the AUROC score dropped and the PSTH shape could not be captured. Therefore, soma distance and synapse activation time point need to be included at high resolution (50µm spatial bins, 1ms temporal bins) to accurately capture the input-output computation. In turn, considering the specific cell types of the neurons from which the synaptic inputs originate did not increase the prediction accuracy (**Fig. S3C-E**). Taken together, the AP history, soma-distance dependent spatial distribution of synapses and their temporal activation pattern are necessary and sufficient features to accurately predict APs in the investigated L5PT multi-scale model. Thus, reduced models that are based on these three features, such as the GLM described here, provide an analytically tractable and interpretable description of the input-output computation that a L5PT with this morphology and these biophysical properties performs upon sensory stimulation.

### Input-output computation is robust to morphological and biophysical diversity

How does this input-output computation depend on the morphological and biophysical properties of L5PTs? To address this question, we incorporated the multi-scale model configurations of all multi-compartmental models with morphologically and biophysically diverse dendritic trees and applied the reduction strategy to each of them. The reduction revealed that the shape of the spatiotemporal filters, the shape of the penalty and nonlinearity are qualitatively similar across all multi-compartmental models despite their morphological and biophysical diversity (**Fig. 3A**): All L5PTs count active synapses, strongly weighting proximal input that occurs within the characteristic EPSP/IPSP-like time windows, with a similar AP-history dependent penalty. Each reduced model achieved a high accuracy before and after the sensory stimulus (**Fig. 3B**), and highest accuracies during the peak response (AUROC median/min/max: 0.97/0.93/0.998). Thus, this input-output computation is robust across morphologically and biophysically diverse L5PTs.

**Figure 3:**
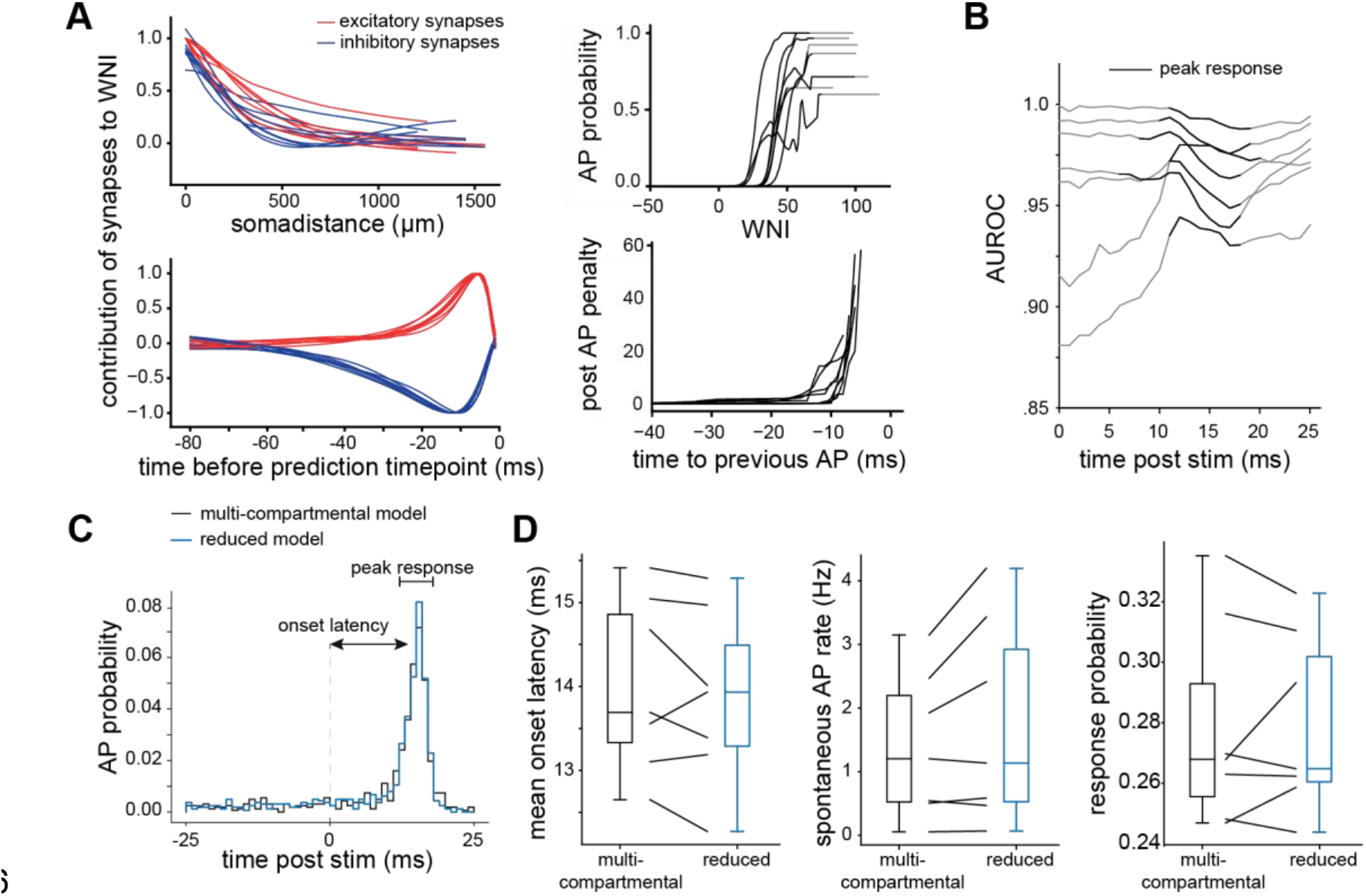
Input-output computation is robust to morphological and biophysical diversity. **A:** Reduced models inferred on the different multi-compartmental models are qualitatively similar, with similar temporal and spatial filters, nonlinearity and post AP penalty. **B:** All models have high AUROC scores, specifically during the sensory-evoked (peak) response. **C:** We quantify latency, spontaneous AP rate (before the stimulus) and response probability for each pair of multi-compartmental and reduced model. **D:** Comparing response properties between multi-compartmental and corresponding reduced model shows close match.

Notably, while the properties of the reduced models are qualitatively similar, they are not identical. To what degree do these small quantitative variations in the reduced model capture differences of the input-output computations across L5PTs? We compared the predicted PSTHs between the multi-scale configurations of each multi-compartmental model with the respective reduced model and found that the slight differences between their shapes are captured well by the reduced model (**Fig. S4**). To quantify this similarity, we compared the time to maximum response (latency), AP rate before the stimulus, and response probability between multi-compartmental models and corresponding reduced models (**Fig. 3C**). Each reduced model generated responses which matched those of the corresponding multi-compartmental models in all of these properties, while preserving the considerable variation across them (**Fig. 3D**). Thus, the small variability in the reduced models account for differences in the input-output computation between the multi-compartmental models.

### Reduced models explain origins of receptive field variability

How can the large variability in L5PT responses arise from such small variability in input-output computations? We repeated the simulations of all multi-scale model configurations, this time with the reduced models instead of the multi-compartmental models. We compared the response probability (i.e. the probability of >=1 AP 0-25ms post passive whisker deflection) between reduced and multi-compartmental models for each whisker, network embedding, and L5PT model, which were highly correlated (**Fig. 4A**, Pearson correlation coefficient = 0.97). Consequently, also the receptive fields were virtually identical (Pearson R between receptive fields 0.95±0.1, **Fig. 4B and C**). On the single trial level, the accuracy of the reduced model was 90.87% ± 3.93% (min 84.96%, max 96.74%). Thus, the reduced models accurately capture the responses of the multi-compartmental model, independent of network embedding, biophysical properties, and morphology and capture cell-to-cell and trial-to-trial response variability.

**Figure 4:**
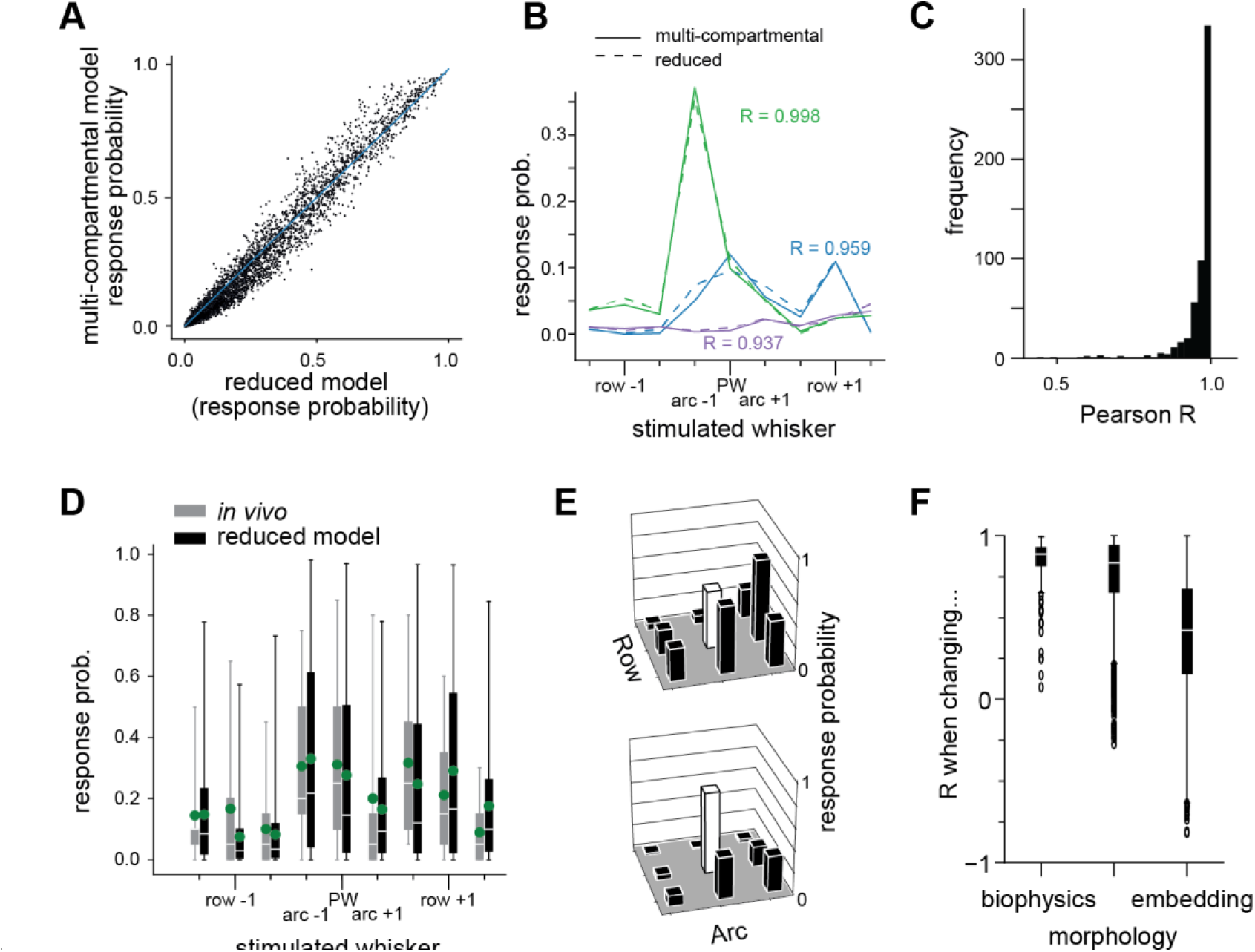
Reduced models explain origins of receptive field variability. **A:** Comparison of responses of 7 different multi-compartmental models and their corresponding reduced models to 9 different whisker stimuli (PW and 8 SW) in 81 different network embedding locations. Response probability is the probability that one or more APs are generated 0-25ms after the sensory stimulus. **B:** Comparison of exemplary receptive field shapes shows close match between biophysically detailed and reduced model. **C:** Quantification of receptive field similarity for all cell positions and biophysically detailed models. **D:** Comparison between *in vivo* and reduced model responses to 9 different whisker stimuli (PW and 8 SW). **E:** Exemplary receptive fields of reduced models. **F:** Influence of biophysics, morphology and cell position on receptive field shape, quantified by computing the correlation coefficient between receptive fields if one of these properties is changed.

How well do the reduced model responses match *in vivo* data? We compared response probabilities to each whisker stimulus (PW and 8 SWs) between *in vivo* data and *in silico* predictions (**Fig. 4D**). The distribution of response probabilities closely match for any whisker (p values ranging between 0.09 and 0.89) and the means of these distributions, i.e., the ‘mean receptive field’, is significantly correlated (R value 0.77, p = 0.015). The simulated receptive fields even contain matches to the extreme cases observed *in vivo* -weaker response to the PW than to a SW, selective response to few whiskers, unselective response to virtually all whiskers (**Fig. 4E**). Thus, the reduced models predict means, variance and outliers consistent with those observed *in vivo*, despite the small variability in input-output computations.

Could the variable responses arise from cell-to-cell variability in input? We investigated the extent of receptive field variation across different morphologies, biophysical parameters, and network embeddings within the barrel column (**Fig. 4F**) by computing the pairwise correlation between them. We found a change in embedding, and therefore network input, to have the strongest effect (i.e. leads to the largest drop in the Pearson correlation coefficient), followed by the neuron’s morphology and biophysical properties. The models thereby indicate that variations in network input are the primary determinant of receptive field variability from cell to cell, whereas variations in cellular properties such as morphology and biophysics may play a minor role. Thus, the reduced model predicts that L5PTs of diverse morphologies and biophysical properties perform essentially the same input-output computation upon sensory stimulation, and the variability in their responses is determined by the input they receive from the network.

### Reduced models explain contribution of input pathways to responses

Which presynaptic neuron populations underlie the variable AP responses? While so far we only distinguished between excitatory and inhibitory cells, the network model reports the origin for each active synapse, for both thalamocortical (i.e., VPM) and excitatory intracortical cell types: pyramidal neurons in L2/3/4 (L2PY, L3PY, L4PY), spiny neurons in L4 (L4SP), intratelencephalic neurons in L5 (L5IT), L5PT, corticocortical neurons at the L5/6 border (L6CC) and in deep L6 (L6INV), and corticothalamic neurons in L6 (L6CT). To dissect, how these presynaptic populations contribute to AP output, we utilized the reduced models and calculated the WNI separately for each cell type across whisker stimuli and network embeddings.

First, we analyzed L5PTs embedded at the center of the column receiving a PW (i.e. somatotopically aligned) stimulus. The analysis shows that presynaptic neurons contribute in two ways to APs, either by their spontaneous activity (‘baseline WNI’), or by their increase in activity upon sensory stimulation (‘sensory-evoked WNI’). Preceding the stimulus, predominantly other L5PT neurons contribute to the baseline WNI (see **Fig. 5A** baseline), due to their high spontaneous firing rate [8,13]. To isolate the effect of the sensory-evoked WNI, we subtracted the baseline (**Fig. 5B**). After the stimulus, VPM provides the first sensory-evoked contribution to the WNI, followed by L6CC which is the first intracortical cell type that responds reliably to VPM input [10]. For a SW stimulus (**Fig. 5C**), this is different: VPM contributes only marginally to AP output, while L6CC neurons provide the main drive. Thus, the reduced model predicts that depending on the stimulated whisker, different thalamocortical and intracortical pathways -primarily VPM and L6CC -contribute to the sensory-evoked response.

**Figure 5:**
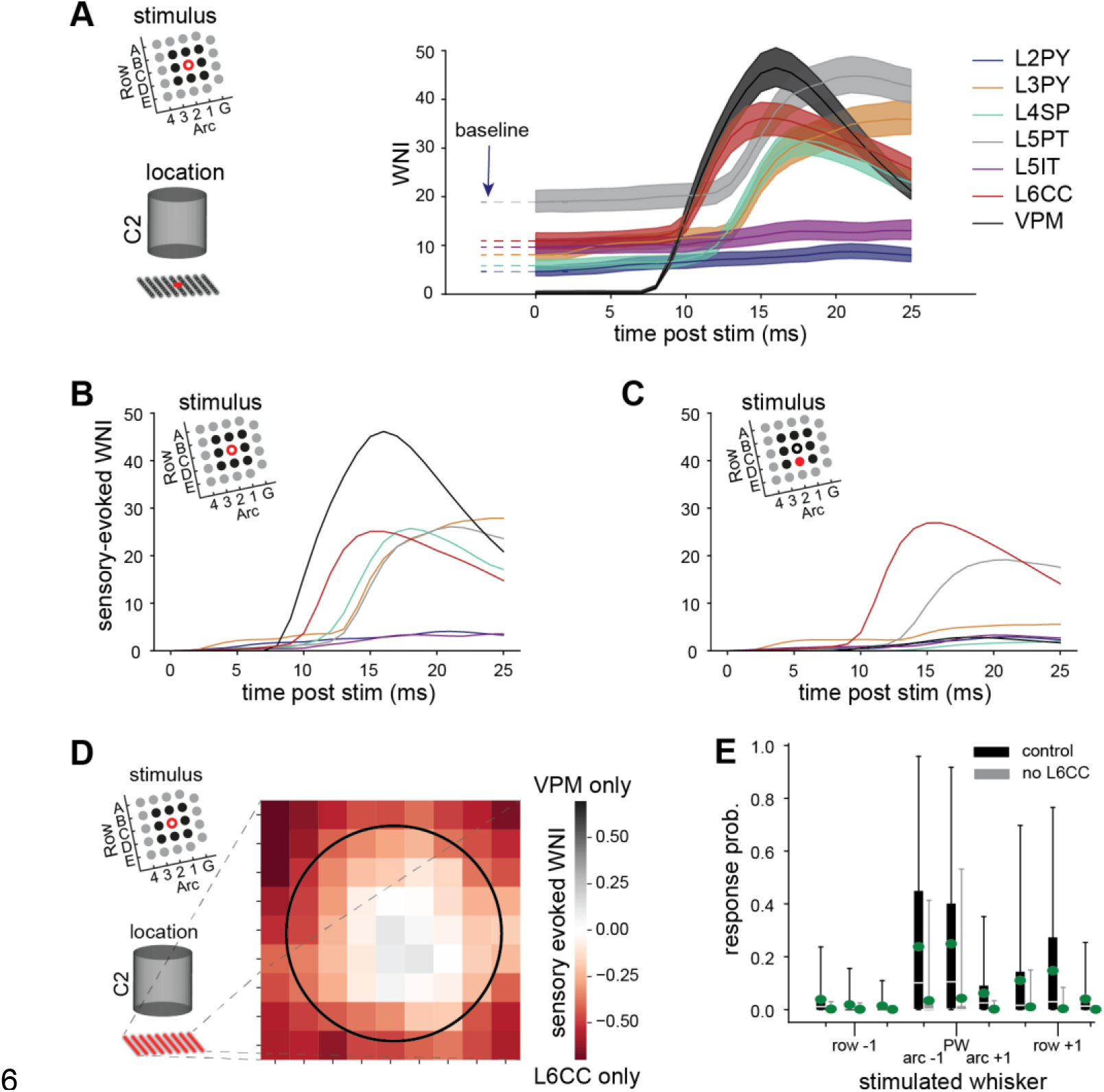
Reduced models explain contribution of input pathways to sensory-evoked responses. **A:** Absolute contribution of presynaptic populations to WNI following a PW stimulus to a model located at the center of the C2 column: pyramidal neurons in L2/3 (L2PY, L3PY), spiny neurons in L4 (L4SP), intratelencephalic neurons in L5 (L5IT), L5PT, corticocortical neurons at the L5/6 border (L6CC), and relay cells in the ventral posterior medial nucleus of thalamus (VPM). **B:** Sensory-evoked contribution (i.e. absolute contribution minus baseline) of presynaptic populations to WNI following a PW stimulus. **C:** Sensory-evoked contribution of presynaptic populations to WNI following a SW stimulus. **D:** Contribution of the main input pathways – VPM and L6CC – depending on the soma location of the L5PT model in a 9x9 grid across the C2 column for a PW stimulus. The black circle denotes the C2 column border. **E:** Comparison between model responses to 9 different whisker stimuli (PW and 8 SW) under control conditions and when removing sensory-evoked input from L6CC. Removing evoked L6CC activity attenuates responses, in particular to surround whiskers.

Could variability in the input from these two pathways underlie the heterogeneous receptive fields of L5PTs? We computed, which fraction of the WNI is provided by VPM versus L6CC depending on the network embedding for a PW stimulus (**Fig. 5D**). This revealed that input from VPM dominates at the center of the barrel column. Towards the column border, VPM contribution decreases whereas the relative input from L6CC increases. L5PTs therefore receive heterogeneous input from these pathways depending on their location. Thus, L6CC provides strong input both for PW and SW stimuli. Inactivating L6CC – a manipulation which we have previously performed experimentally [10] – should therefore strongly reduce SW responses (as L6CC input is the main driver) and reduce PW responses (as L6CC is a strong contributor in this condition), resulting in narrow receptive fields. We repeated the simulations with the reduced model in the absence of sensory-evoked L6CC input for all whiskers. Indeed, we see a major reduction in the overall response, and a narrowing of receptive fields across all model configurations and network embeddings (**Fig. 5E**). Thus, the reduced model predicts that fast responses of L5PTs and broad receptive fields require input from L6CCs, consistent with experimental observations [10].

## Discussion

In this paper we sought to reveal the input-output computations that drive sensory-evoked responses in L5PTs. By deriving analytically tractable, interpretable models from biologically detailed multi-scale models, we identify three features that are sufficient to explain the *in vivo* responses and receptive fields of these neurons. By generating a GLM, which is based on these features, we demonstrate that the input-output computation is robust with respect to variations in morphology and biophysical properties of the dendrites. Diversity in dendritic properties is hence predicted to have only a minor contribution to trial-to-trial and cell-to-cell variability of sensory-evoked responses and receptive fields. Instead, we show that variations in input is likely to account for the *in vivo* observed response variability. The GLM also predicts how manipulations of presynaptic populations impact L5PT responses *in vivo*. Thus, the GLM can account for the mechanistic origins of L5PT responses that have been observed empirically.

A popular choice for modelling how neurons respond to network input are leaky integrate-and-fire models, which are computationally efficient but have no representation of the constituents of a biological neuron, and a priori *assume* the input-output computation of the neuron. In contrast, the approach presented here starts with morphologically and biophysically diverse detailed neuron models, and incorporates electrophysiological data to capture the characteristic dendritic and somatic properties. Next, the input-output computation is *derived* by exposing these detailed models to well-constrained spatiotemporal synaptic input patterns of the investigated experimental condition. The input-output computations are captured in reduced models which inherit the properties of the detailed models; in other words, the reduced models are equivalent to the detailed models thereby representing a biological neuron. We find reduced models that are computationally similarly efficient to leaky integrate-and fire models, but which are based on three features that we show are necessary and sufficient to capture the input-output computation. As they are derived from morphologically and biophysically diverse multi-compartmental models, they account for biological variability in dendritic properties. Thereby, they set the stage for simulations of networks with high biological realism at similar computational cost to integrate-and-fire networks.

We want to emphasize that the reduced models are derived for one specific set of experimental conditions – passive deflections of single whiskers in anaesthetized animals. While we find that the input-output computation of L5PTs under this condition can be captured in very simple models, this is likely not the case in general. Previous studies have shown how to convert multi-compartmental models into deep artificial neural networks [14], simplified conductance based models [15,16] or stacks of linear-nonlinear units [17], which maintain high accuracy throughout a wide range of input conditions. All of these models are highly complex -for example Beniaguev et al. (2021) find that a 7-layer convolutional network is necessary to capture the immense computational power of a single neuron. Here, we have explored an orthogonal approach: by limiting the synaptic input conditions to those present in the specific investigated experimental condition and thereby compromising on the generalizability of the model to other input scenarios, we can derive highly accurate yet interpretable and analytically tractable models that explain how *in vivo* observed responses arise from cellular and circuit properties.

Generating interpretable reduced models for another condition would require first generating a multi-scale model that accounts for the spatiotemporal input pattern and response variability of this condition, and searching for features that can predict the responses. These features may not fit a GLM structure. For example, in awake animals, specifically the occurrence of sensory-evoked high-frequency bursts is increased in L5PTs [18]. These bursts likely originate from non-sensory information streams impinging onto the distal dendrites, which thereby facilitate the occurrence of dendritic calcium APs [19], for which the current GLM models cannot account (**Fig. S4**). While such scenarios are extremely rare under the investigated condition, they are more frequent for awake conditions. Revisiting trials which are misclassified by the current reduced model and determining which additional features are needed to accurately predict the responses would thereby result in a new minimal description of the input-output computation performed by neurons under this new experimental condition.

While presently we rely on a multi-scale model as the basis for our reduction approach, in the coming years, large-scale voltage-imaging and connectome data at electron microscopic resolution may become increasingly available. Such experimental approaches could provide data at a similar level of detail to the multi-scale model. The reduction approach we present here could equally be applied to such future experimental data, to understand how the observed cellular activity can arise from cellular and network properties. Thus, our approach provides a roadmap for revealing input-output computations and their cellular and network implementation across different *in vivo* conditions.

## Methods

## Data availability statement

Code for reduced model inference is provided here: https://github.com/mpinb/BastFruengelEtAl/blob/main/BastFruengelEtAl.ipynb

Training data for the reduced models (synapse activation times and spike times) is provided here for review purposes and will be made available in a public repository upon publication: https://web.tresorit.com/l/kjlLA#7qSe1amSVnOCsDGkft-Lxg

### Extracellular recordings

The electrophyisological data used in this study has been reported previously [8]. Briefly, Wistar rats (P33-P70, m) were anesthetized using urethane and responses to single whisker stimuli (applied with a piezo manipulator) of the PW and SWs were recorded via a cell attached pipette. The neurons were filled with biocytin and morphologically reconstructed to identify their cell types. The PW was determined by identifying the barrel column in which the soma of the recorded L5PT was located – i.e., the PW is not necessarily the one that evoked the strongest response. The *in vivo* data used in this study comprised exclusively L5PTs located in the D2 column.

### Morphological reconstructions

Neuronal structures were extracted from image stacks using a previously reported automated tracing software [20]. For reconstruction of biocytin labeled neurons, images were acquired using a confocal laser scanning system (Leica Application Suite Advanced Fluorescence SP5; Leica Microsystems). 3D image stacks of up to 2.5mm × 2.5mm × 0.05 mm were acquired at 0.092 × 0.092 × 0.5μm per voxel (63x magnification, NA 1.3). Image stacks were acquired for each of 45-48 consecutive 50μm thick tangential brain slices that range from the pial surface to the white matter. Manual proof-editing of individual sections, and automated alignment across sections were performed using custom-designed software [21]. Pia, barrel and white matter outlines were manually drawn on low-resolution images (4x magnification dry objective). Using these anatomical reference structures, all reconstructed morphologies were registered to a standardized 3D reference frame of rat vS1 [22].

### Multi-compartmental models

We selected 5 L5PT reconstructions that are representative of the morphological variability of this cell type. Multi-compartmental models were generated for these morphologies as described previously [10,12]. Briefly, a simplified axon morphology was attached to the soma of the reconstructed L5PT dendrite morphology [23]. The axon consisted of an axon hillock with a diameter tapering from 3μm to 1.75μm over a length of 20μm, an axon initial segment of 30μm length and diameter tapering from 1.75μm to 1μm diameter, and 1 mm of myelinated axon (diameter of 1μm). Next, a multi-objective evolutionary algorithm was used to find parameters for the passive leak conductance and the density of Hodgkin-Huxley type ion channels on soma, basal dendrite, apical dendrite and axon initial segment, such that the neuron model is able to reproduce characteristic electrophysiological responses to somatic and dendritic current injections of L5PTs within the experimentally observed variability, including back-propagating APs, calcium APs, and AP responses to prolonged somatic current injections [12]. We augmented the original biophysical model of L5PTs [10,12] with two ion channel parameters as previously described [24]: in accordance with a previous report [25], the density of the fast non-inactivating potassium channels (Kv3.1) was allowed to linearly decrease with soma distance until it reaches a minimum density (i.e., the slope and minimum density are two additional parameters, see [24]). The diameter of the apical dendrites was optimized by a scaling factor between 0.3 and 3. We incorporated the IBEA algorithm [26] for optimization. The optimization was terminated if there was no progress or when acceptable models had been found. We repeated the optimization process several times. From each independent run, we selected one model for which the maximal deviation from the experimental mean in units of standard deviation across all objectives was minimal (0.9-1.9 mean STDs across objectives).

### Network embeddings

Cell type-specific thalamocortical (from the ventral posteromedial thalamic nucleus (VPM) which is the primary thalamic nucleus of the whisker) and intracortical (from excitatory and inhibitory cells in vS1) connections are derived from an anatomically realistic circuit model of rat vS1 [9], a procedure, which has been described in detail previously [10]. We embedded the dendrite morphologies selected for multi-compartmental modelling in the network model at 81 locations within the cortical barrel column representing the C2 whisker, which is located approximately in the center of vS1. For all *in silico* data presented in this study, the PW is hence the C2 whisker. Thus, the morphologies of the *in vivo* recorded L5PTs were registered to the C2 column, while preserving their *in vivo* soma depths and laminar dendrite distributions that was observed empirically in the D2 column (Egger, 2012). The locations were the column center, and equally spaced grid with a distance of 50μm between adjacent somata. For each of the 81 locations, we estimated the location of presynaptic neurons in VPM and vS1 that provide input to the respective L5PT (**Fig. 1B**) and where along the L5PT dendrite they synapse (**Fig. 1C**).

### Synapse models

Synapse models and synaptic parameters (rise and decay times, release probabilities, reversal potentials) were reported previously [10]. Briefly, conductance-based synapses were modeled with a double-exponential time course. Excitatory synapses contained both AMPA receptors (AMPARs) and NMDARs in 1:1 ratio. Inhibitory synapses contained GABA ARs. The peak conductance of excitatory synapses from different presynaptic cell types was determined by assigning the same peak conductance to all synapses of the same cell type, activating all connections of the same cell type (i.e., all synapses originating from the same presynaptic neurons) one at a time, and comparing parameters of the resulting unitary postsynaptic potential (uPSP) amplitude distribution (mean, median and maximum) for a fixed peak conductance with experimental measurements *in vitro* (IC input [27]) or *in vivo* (TC input [28]). The peak conductance for synaptic inputs from each cell type was systematically varied until the squared differences between parameters of the *in silico* and *in vitro/in vivo* uPSP amplitude distributions were minimized (i.e., the mean, median and maximum of the distributions were used, and mean and median were weighted twice relative to the maximum). This procedure was repeated for each multi-compartmental model using the connectivity model for the location in the center of the C2 column. The peak conductance at inhibitory synapses was fixed at 1nS [13].

### Synaptic input patterns

Synaptic input patterns to the L5PT model were estimated as described previously [10]. Briefly, we first chose the whisker stimulus. Then presynaptic neurons were grouped by their cell type and the column in which they are located. For each column, the relative position to the stimulated whisker was determined. Neurons in this column were activated based on the electrophysiologically recorded responses for this cell type and relative position. Each AP in a presynaptic neuron is registered at all synapses between the presynaptic neuron and the L5PT model without delay and may cause a conductance change, depending on the release probability of the synapse. Depending on the network embedding, neurons receive different ratios of excitatory and inhibitory inputs, which is not guaranteed to maintain functional E/I balance [29]. We therefore introduced a scaling factor for evoked inhibitory input strength, which we constrained based on the empirically observed PW response probabilities. The scaling factors ranged between 0.79 and 1.56 depending on the multi-compartmental model.

### Multi-compartmental model simulation data

Simulations were performed using Python 2.7, dask [30] and NEURON 7.4 [31]. We simulated 1000 trials for each of the 9 multi-compartmental models and each of the 81 locations and 9 whisker stimuli (PW, all 8 SWs). Additionally, the database contained 81000 trials of one 2^nd^ SW (E2), which was however not incorporated in the analysis. This results in 810000 simulation trials per multi-compartmental model, which reflects the effect of biophysics, morphology, network embedding and stimulus on sensory-evoked responses.

### Reduced model inference

We performed the reduced model inference for each multi-compartmental model separately. We split the multi-compartmental model dataset for the respective model into a training and test dataset (split ratio: 70% to 30%, respectively). We excluded trials of the recent AP category (in which there was an AP in the last 50ms). Spatiotemporal filters were parametrized with a raised cosine basis function ([32], see **Fig. S6**). Parameters determining the shape of the spatial and temporal kernels of the reduced model were adjusted by a gradient-free optimization method (COBYLA, implemented in SciPy: [33]) such that the area under the receiver operating characteristic curve (AUROC) is maximized. Using these spatiotemporal filters, we weight all active synapses preceding the prediction time point depending on their individual soma distance, relative activation time point and synapse type (excitatory or inhibitory). This results in a single, scalar value which is predictive for spiking. In the following, we refer to this value as weighted net input (WNI), as it reflects the net amount of excitation the neuron received. The WNI is computed as follows:

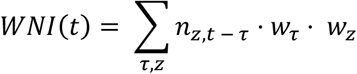

where *t* is the prediction timepoint, *τ* ϵ [0, 80ms] is the time before the prediction timepoint, *z* ϵ [0, 1300μm] is distance of the synapse from the soma, and *n*_*z,t* − *τ*_ is the number of active synapses at a given time and soma distance. *w*_*τ*_ and *w*_*z*_ are the temporal and spatial filter, respectively.

These filters were constructed as a linear sum from basis functions *f*_*i*_ and *g*_*j*_:

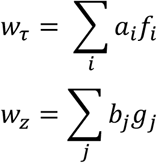

where *a*_*i*_ and *b*_*j*_ are free parameters which were subjected to the optimizer. We chose a raised cosine basis for *f*_*i*_ and *g*_*j*_:

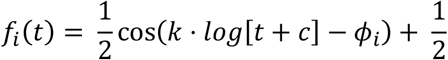

for t such that k·log(t + c) ϵ [φi – π, φi + π] and 0 elsewhere. Values used were k = 3, c = 5, and φ ϵ [3, 12]. Analogously, g is also a raised cosine function:

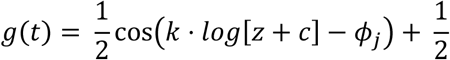

for z such that k·log(z + c) ϵ [φi – π, φi + π] and 0 elsewhere. Values used were k = 2, c = 1, and φ ϵ [1, 11].

In order to calculate the spiking probability for a given WNI value, we used these filters to calculate the WNI for all multi-compartmental model simulation trials from the training dataset at the inference time point, and recorded whether a AP occurred at this time point or not. The trials were then binned by WNI. The AP probability for each bin corresponds to the proportion of trials that produced an output AP in the multi-compartmental model simulation. Bins with few data points were combined to ensure a minimum of ten data points per bin. Linear interpolation was used to find the spiking probability corresponding to any WNI value, and WNI values greater/smaller than values seen in the biophysical model simulation were assigned the highest/lowest spiking probability respectively. We hereafter refer to this as the “nonlinearity” function. In order to estimate the effect of recent APs on spiking probability, WNI values were plotted against the time since the previous AP (**Fig. 2C**), and a boundary was drawn to best separate spiking from non-spiking trials as follows. We dropped points in the lowest 5% of all WNI values (to remove the effect of outliers) and then drew a boundary along the minimum remaining spiking WNI values. This boundary was normalized such that it is zero for time to previous AP → ∞ by subtracting the respective offset. We found this two-step procedure (first estimate spatiotemporal filters based on a dataset in which recent APs are filtered out, second determine the post AP penalty based on this filters) to be more robust than a joint inference of both. WNI with penalty applied maintains a high AUROC over all stimulus periods (**Fig. 2F**), while the uncorrected AUROC drops after the sensory stimulus.

### Simulations with reduced models

APs were generated based on the synaptic input reflecting whisker stimuli generated above. AP output was generated iteratively for one millisecond time bin at a time. For each time point, we computed the WNI by applying the temporal and spatial filter to the synaptic input of the preceding 80ms. The WNI was transformed to an AP probability based on the nonlinearity derived above. APs were randomly sampled based on this probability. If an AP occurred, the WNI of consecutive time points was updated based on the post AP penalty.

### Analysis

‘AP probability’ is used to denote the probability that an AP is evoked within a 1ms time bin. ‘Response probability’ denotes the probability that one or more APs are generated in the 25ms following a whisker stimulus.

## Acknowledgements

***Funding*** was provided by the Max Planck Institute for Neurobiology of Behavior – caesar (MO), European Research Council grants 633428 and 101069192 (MO), Deutsche Forschungsgemeinschaft grants SFB 1089 and SPP 2041 (MO), German Federal Ministry of Education and Research grant 01IS18052 (MO), and Neuroscience Network North Rhine-Westphalia grant iBehave (MO).

## Author contributions

AB and MO conceived and designed the study. AB developed the model reduction approach. AB and RF performed simulations and analyzed the data. CK contributed *in vivo* data. AB and RF wrote the paper with MO.

## References

1. Chen X, Leischner U, Rochefort NL, Nelken I, Konnerth A. Functional mapping of single spines in cortical neurons in vivo. Nature. 2011;475: 501–505. doi:10.1038/nature10193

2. Jia H, Varga Z, Sakmann B, Konnerth A. Linear integration of spine Ca ^2+^ signals in layer 4 cortical neurons in vivo. Proc Natl Acad Sci USA. 2014;111: 9277–9282. doi:10.1073/pnas.1408525111

3. Rochefort NL, Konnerth A. Dendritic spines: from structure to in vivo function. EMBO Reports. 2012;13: 699–708. doi:10.1038/embor.2012.102

4. Scholl B, Thomas CI, Ryan MA, Kamasawa N, Fitzpatrick D. Cortical response selectivity derives from strength in numbers of synapses. Nature. 2021;590: 111–114. doi:10.1038/s41586-020-03044-3

5. Varga Z, Jia H, Sakmann B, Konnerth A. Dendritic coding of multiple sensory inputs in single cortical neurons in vivo. Proc Natl Acad Sci USA. 2011;108: 15420–15425. doi:10.1073/pnas.1112355108

6. Woolsey TA, Van Der Loos H. The structural organization of layer IV in the somatosensory region (S I) of mouse cerebral cortex. Brain Research. 1970;17: 205–242. doi:10.1016/0006-8993(70)90079-X

7. Harris KD, Shepherd GMG. The neocortical circuit: themes and variations. Nat Neurosci. 2015;18: 170–181. doi:10.1038/nn.3917

8. de Kock CPJ, Bruno RM, Spors H, Sakmann B. Layer- and cell-type-specific suprathreshold stimulus representation in rat primary somatosensory cortex. The Journal of physiology. 2007;581: 139–154. doi:10.1113/jphysiol.2006.124321

9. Egger R, Dercksen VJ, Udvary D, Hege H-C, Oberlaender M. Generation of dense statistical connectomes from sparse morphological data. Front Neuroanat. 2014;8: 129. doi:10.3389/fnana.2014.00129

10. Egger R, Narayanan RT, Guest JM, Bast A, Udvary D, Messore LF, et al. Cortical Output Is Gated by Horizontally Projecting Neurons in the Deep Layers. Neuron. 2020;105: 122–1378. doi:10.1016/j.neuron.2019.10.011

11. Udvary D, Harth P, Macke JH, Hege H-C, de Kock CPJ, Sakmann B, et al. The impact of neuron morphology on cortical network architecture. Cell Reports. 2022;39: 110677. doi:10.1016/j.celrep.2022.110677

12. Hay E, Hill S, Schürmann F, Markram H, Segev I. Models of neocortical layer 5b pyramidal cells capturing a wide range of dendritic and perisomatic active properties. PLoS computational biology. 2011;7: e1002107. doi:10.1371/journal.pcbi.1002107

13. Hay E, Segev I. Dendritic Excitability and Gain Control in Recurrent Cortical Microcircuits. Cerebral cortex (New York, NY : 1991). 2015;25: 3561–3571. doi:10.1093/cercor/bhu200

14. Beniaguev D, Segev I, London M. Single cortical neurons as deep artificial neural networks. Neuron. 2021;109: 2727–27393. doi:10.1016/j.neuron.2021.07.002

15. Amsalem O, Eyal G, Rogozinski N, Gevaert M, Kumbhar P, Schürmann F, et al. An efficient analytical reduction of detailed nonlinear neuron models. Nat Commun. 2020;11: 288. doi:10.1038/s41467-019-13932-6

16. Pagkalos M, Chavlis S, Poirazi P. Introducing the Dendrify framework for incorporating dendrites to spiking neural networks. Nat Commun. 2023;14: 131. doi:10.1038/s41467-022-35747-8

17. Ujfalussy BB, Makara JK, Lengyel M, Branco T. Global and Multiplexed Dendritic Computations under In Vivo-like Conditions. Neuron. 2018;100: 579–5925. doi:10.1016/j.neuron.2018.08.032

18. De Kock CPJ, Pie J, Pieneman AW, Mease RA, Bast A, Guest JM, et al. High-frequency burst spiking in layer 5 thick-tufted pyramids of rat primary somatosensory cortex encodes exploratory touch. Commun Biol. 2021;4: 709. doi:10.1038/s42003-021-02241-8

19. Bast A, Guest JM, Fruengel R, Narayanan RT, De Kock CPJ, Oberlaender M. Thalamus drives active dendritic computations in cortex. Neuroscience; 2021 Oct. doi:10.1101/2021.10.21.465325

20. Oberlaender M. Transmitted light brightfield mosaic microscopy for three-dimensional tracing of single neuron morphology. J Biomed Opt. 2007;12: 064029. doi:10.1117/1.2815693

21. Dercksen VJ, Hege H-C, Oberlaender M. The Filament Editor: An Interactive Software Environment for Visualization, Proof-Editing and Analysis of 3D Neuron Morphology. Neuroinform. 2014;12: 325–339. doi:10.1007/s12021-013-9213-2

22. Egger R, Narayanan RT, Helmstaedter M, De Kock CPJ, Oberlaender M. 3D Reconstruction and Standardization of the Rat Vibrissal Cortex for Precise Registration of Single Neuron Morphology. Briggman K, editor. PLoS Comput Biol. 2012;8: e1002837. doi:10.1371/journal.pcbi.1002837

23. Hay E, Schürmann F, Markram H, Segev I. Preserving axosomatic spiking features despite diverse dendritic morphology. Journal of Neurophysiology. 2013;109: 2972–2981. doi:10.1152/jn.00048.2013

24. Bast A, Oberlaender M. Ion channel distributions in cortical neurons are optimized for energy-efficient active dendritic computations. Neuroscience; 2021 Dec. doi:10.1101/2021.12.11.472235

25. Schaefer AT, Helmstaedter M, Schmitt AC, Bar-Yehuda D, Almog M, Ben-Porat H, et al. Dendritic voltage-gated K ^+^ conductance gradient in pyramidal neurones of neocortical layer 5B from rats: K ^+^ conductance gradient in pyramidal neurones. The Journal of Physiology. 2007;579: 737–752. doi:10.1113/jphysiol.2006.122564

26. van Geit W, Gevaert M, Chindemi G, Rössert C, Courcol J-D, Muller EB, et al. BluePyOpt: Leveraging Open Source Software and Cloud Infrastructure to Optimise Model Parameters in Neuroscience. Frontiers in neuroinformatics. 2016;10: 17. doi:10.3389/fninf.2016.00017

27. Schnepel P, Kumar A, Zohar M, Aertsen A, Boucsein C. Physiology and Impact of Horizontal Connections in Rat Neocortex. Cereb Cortex. 2015;25: 3818–3835. doi:10.1093/cercor/bhu265

28. Constantinople CM, Bruno RM. Deep cortical layers are activated directly by thalamus. Science (New York, NY). 2013;340: 1591–1594. doi:10.1126/science.1236425

29. Landau ID, Egger R, Dercksen VJ, Oberlaender M, Sompolinsky H. The Impact of Structural Heterogeneity on Excitation-Inhibition Balance in Cortical Networks. Neuron. 2016;92: 1106–1121. doi:10.1016/j.neuron.2016.10.027

30. Dask Development Team. Dask: Library for dynamic task scheduling. 2016. Available: https://dask.org

31. Hines ML, Carnevale NT. The NEURON simulation environment. Neural computation. 1997;9: 1179–1209. doi:10.1162/neco.1997.9.6.1179

32. Weber AI, Pillow JW. Capturing the dynamical repertoire of single neurons with generalized linear models. 2016. Available: http://arxiv.org/pdf/1602.07389v3

33. Virtanen P, Gommers R, Oliphant TE, Haberland M, Reddy T, Cournapeau D, et al. SciPy 1.0: fundamental algorithms for scientific computing in Python. Nature methods. 2020;17: 261–272. doi:10.1038/s41592-019-0686-2

